# Direct purification of CRISPR/Cas ribonucleoprotein from *E. coli* in a single step

**DOI:** 10.1101/377366

**Authors:** Siyu Lin, Jie Qiao, Lixin Ma, Yi Liu

## Abstract

CRISPR/Cas ribonucleoprotein (RNP) complexes have been recently used as promising biological tools with plenty of applications, however, there are by far no efficient methods to prepare them at large scale and low cost. Here, we present a simple method to directly produce and purify Cas RNP, including the widely used Cas9 and Cas12a nuclease, from *E.coli* in a single step using an ultra-high-affinity CL7/Im7 purification system. The prepared Cas RNP shows high stability, solid nuclease activity in vitro, and profound genome editing efficiency in vivo. Our method is convenient, cost-effective, and applicable to prepare other CRISPR associated nucleases.

## Introduction

Clustered regularly interspaced short palindromic repeats (CRISPR)/CRISPR associated-protein (Cas) genome-editing systems, including the widely used *Streptococcus pyogenes* Cas9 (*Sp*Cas9) (Jinek et al. 2012; Koonin et al. 2017) and *Francisella novicida* Cas12a (previously named FnCpf1) nuclease (Fonfara et al. 2016; Zetsche et al. 2015), have been extensively applied to biomedical area with great promises to revolutionize the treatment of genetic diseases (Cho et al. 2013; Scott and Zhang 2017). Gene therapy with adeno-associated viruses (AAVs) is currently the most advanced approach for delivering Cas enzymes in vivo (Wu et al.; Yin et al. 2016), though this methodology has limitations such as pre-existing immunity towards AAV in human and small packing size (Schumann et al. 2015). Recently, several researches demonstrated that direct delivery of CRISPR/Cas ribonucleoprotein (RNP) for genome editing in cells and animals has obvious advantages (Gao et al. 2018; Staahl et al. 2017; Yin et al. 2014), such as reduced off-target effects, low toxicity, high editing efficiency, etc. Thus, increasing biopharmaceutical companies are paying greater emphasis on developing Cas RNP-based gene therapy medicines.

The current strategy to construct Cas RNP needs to produce recombinant CRISPR-associated nuclease and the guide RNA (gRNA) individually, followed by assembling them in vitro (Kim et al. 2014). However, the preparation of pure RNP is always time-consuming and expensive (Anders and Jinek 2014). Therefore, the development of an efficient and cost-effective method to produce and purify Cas RNP with high yield and high purity remains a central problem in the field of RNP-based therapeutic gene editing.

In our recent work, we successfully achieved co-expression of Cas9 and guide RNA in *E. coli* to directly prepare Cas9 RNP in vivo (Li et al. 2018). We harnessed two chromatographic steps to purify the Cas9 RNP, including the first affinity purification by Ni-NTA column and the second purification by gel filtration. By the method, it takes us more than three days to prepare pure Cas9 RNP with a relatively low production (~ 10 mg/L), which is still not yet satisfied with large-scale demand in industry. In this report, we utilize an ultra-high-affinity CL7/Im7 purification system (Vassylyeva et al. 2017) developed recently to carry out one-step purification of Cas RNP, including Cas9 and Cas12a, with great purity and improved yield (~ 40 mg/L for Cas9; ~ 50 mg/L for Cas12a). Meanwhile, the whole production process was largely reduced to half a day. The Im7 affinity column used in the work was easily made by ourselves through coupling the recombinant Im7 to agarose beads, and repeatedly regenerated without losing its affinity to the CL7 affinity tag. Together, we establish a valid platform to readily produce CRISPR-associated RNP at large-scale and low cost.

## Materials and methods

### Co-expression vector design and construction

We ordered CL7 gene form Sangon Biotech (Shanghai) Company and assembled it in the N-terminal of Cas9 or Cas12a with a TEV proteinase cleavage site between them. In addition, we preserved the C-terminal 6 x His tag for purifying CL7-tagged fusion enzymes if required. Then, the CL7-Cas9 or CL7-Cas12a fragment was cloned into an engineered cold-shock vector pCold I (Takara) (Qing et al. 2004) by replacing the original cspA promoter with a tac promoter, and by replacing the f1 ori with a T7 promoter (**Fig 1.**). In addition, the vector contains a gRNA transcription element which can be easily replaced by S*alI* digestion. We generated two co-expression plasmids termed pCold CL7-Cas9 and pCold CL7-Cas12a, respectively.

**Fig 1.**
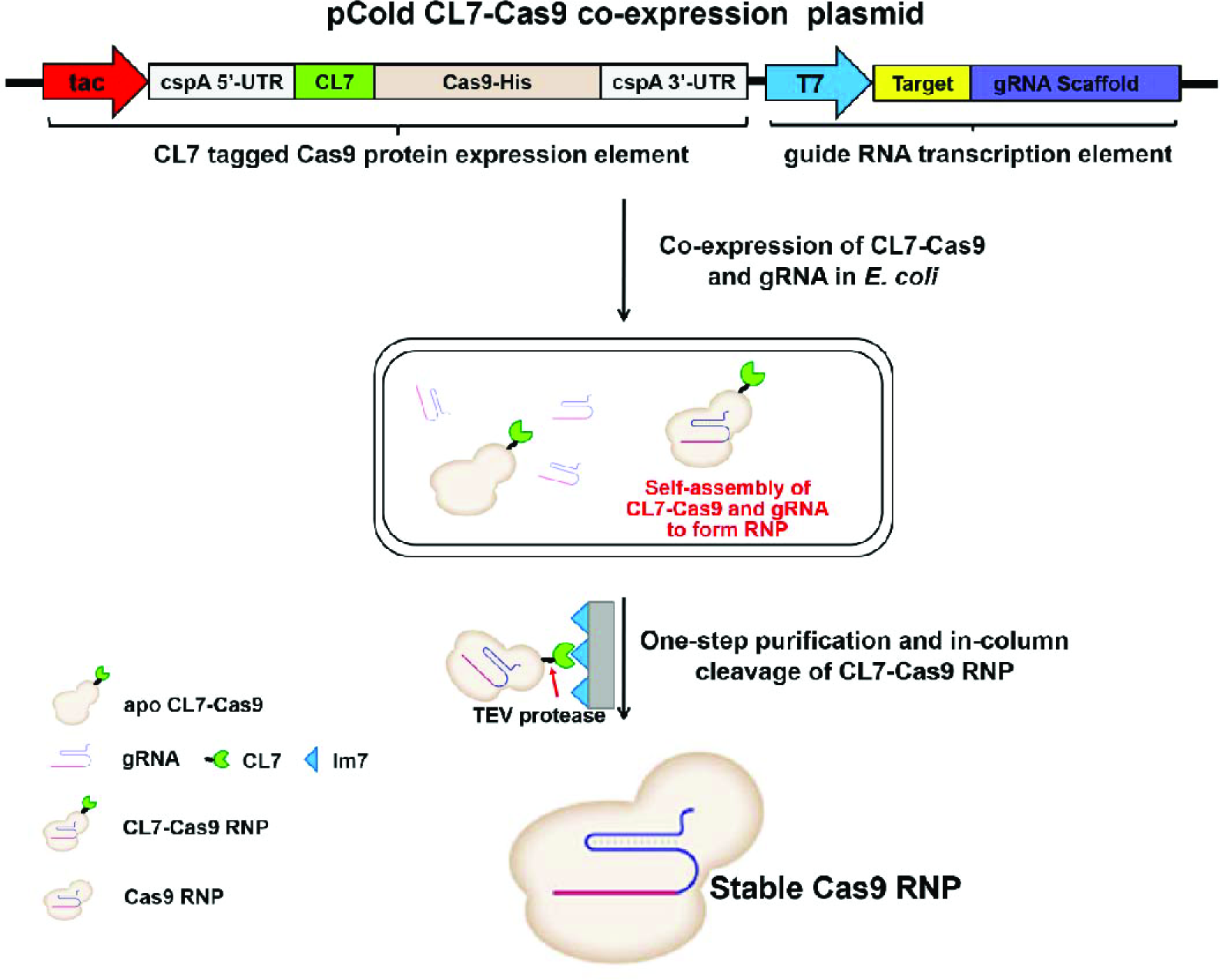
An engineered cold-shock pCold I vector was harnessed to achieve co-expression of CL7-Cas9 and gRNA in *E. coli*. The CL7-Cas9 enzyme and gRNA were self-assembled in vivo to form CL7-Cas9 RNP. The stable Cas9 RNP can be easily obtained by one-step purification and in-column cleavage of the CL7 tag.

### Preparation of Im7 ligated agarose beads

The Im7 gene was synthesized by Sangon Biotech (Shanghai) Company and cloned into an pET23a(+) expression vector (Vassylyeva et al. 2017). The Im7 protein was overexpressed in *E. coli* BL21(DE3) cells and purified according to the protocol. Then, the prepared Im7 was covalently cross-linked to 1 mM iodoacetyl agarose beads purchased from Sigma-Aldrich. After cleavage and elution of a target protein, the CL7-tag, which remained bound to the Im7 agarose beads, was removed under denaturing conditions (6 M Guanidine hydrochloride, Sinopharm Group Co.). The column was then easily regenerated via in-column Im7 refolding.

### Purification of Cas RNP by Im7 column

The co-expression plasmids constructed above were transformed into the *E. coli* Rosetta (DE)3 and the cells were cultured at 37°C under 220 rpm rotate speed. 0.5 mM IPTG was added when the OD600 reached 0.8, and the temperature was decreased to 16°C for expression of Cas RNP. The cells were harvested and lysated in lysis buffer (20 mM Tris-HCl, pH 7.4, 100 mM NaCl). The CL7 tagged Cas9 or Cas12a was loaded onto the Im7 column. After washing two cycles of washing buffer I (20 mM Tris-HCl, pH 7.4, 300 mM NaCl), TEV protease was added to the column for in-column cleavage of CL7 tag for three hours. Next, the highly purified Cas9 or Cas12a protein was eluted with washing buffer II (20 mM Tris-HCl, pH 7.4, 500 mM NaCl). Finally, the Cas RNP was concentrated and stored at −80°C in the storage buffer (20 mM Tris-HCl, pH 7.4, 500 mM NaCl, 20% glycerin).

### In vitro nuclease activity assay

For in vitro endonuclease activity test, the purified Cas RNP was directly applied to digest dsDNA. The digestion was typically carried out in a 10 μl reaction mixture, composed of 1 μl 10 x buffer 3.1 (NEB), 200 ng Cas9 or Cas12a RNP, 300 ng plasmid, at 37°C for 30 minutes followed by termination of the reaction at 80°C for 10 minutes.

### In vivo genome editing efficiency assay

For illustrating in vivo and homology dependent repair (HDR) efficiency, we adopted a BFP-expressing HEK293 reporter cell line according to the reported protocol (DeWitt et al. 2017; Lee et al. 2017). To edit BEP-HEK to GFP, the donor dsDNA and Cas9 RNP were co-delivered by lipofectamine CRISPRMAX (Yu et al. 2016) (ThemoalFisher). The HDR efficiency were calculated from the GFP cell counts/total cell counts (DeWitt et al. 2017; Lee et al. 2017).

## Results

### Direct expression of CL7 tagged Cas RNP in *E. Coli*

To prepare a large amount of Cas RNP, we introduce a N-terminal CL7-tag (Vassylyeva et al. 2017) into the co-expression vector engineered from a commercial pCold I vector (Takara) (Qing et al. 2004) (**Fig. 1**). CL7 is a catalytically inactive variant of Colicin E7 (CE7) DNase with a pretty low Kd (~10^−14^-10^−17^ M) towards its binding partner Immunity Protein 7 (Im7) (Ko et al. 1999). The CL7/Im7 system has been reported recently to facilitate purification of diverse proteins (Vassylyeva et al. 2017). According to the design, the gRNA molecules were abundantly transcribed in vivo by *E. coli’s* own RNA polymerases, while the CL7-Cas9 fusion proteins were largely produced when induced by IPTG. We have proved that the newly synthesized Cas9 and transcribed gRNA molecules would be spontaneously self-assembled in *E. coli* cells to form complete Cas9/gRNA complexes (Li et al. 2018). Likewise, in this case, the CL7 tagged Cas9 would interact with the transcribed gRNA in vivo to form CL7-Cas9 RNP. Introduction of CL7 tag would not only significantly simplify the purification procedures, but also can hugely improve the solubility and yield of target proteins (Vassylyeva et al. 2017). As a result, it brings about five-fold increase in production of Cas9 RNP, compared to the method in absence of CL7 tag. In addition, we applied the method to produce Cas12a nuclease, and observed approximately ten-fold yield compared to the current method that used maltose binding protein (MBP) as fusion tag. More importantly, we found that the Cas9 (or Cas12a) with and without CL7 tag have the same endonuclease activity (**Fig. 3a**), possibly due to CL7 has a small molecular weight (~ 16 kD).

### Purification of pure Cas RNP in a single step

We previously harnessed multiple chromatographic steps to obtain pure Cas9 RNP, therefore, a large number of Cas9 enzymes were lost during the purification procedures. In this work, following an optimized protocol (see SI for details), we successfully achieved purification of Cas RNP in a single step within half a day by the ultra-high-affinity CL7/Im7 system. Comparing to the Cas9 RNP purified by single Ni-NTA column, the purity of Cas9 RNP prepared by Im7 column was significantly improved (**Fig. 2**). More importantly, the Im7 affinity column was easily made (SI) by ourselves through coupling the recombinant Im7 proteins to agarose beads, and repeatedly regenerated without losing its affinity to the CL7 affinity tag, largely reducing the cost of materials.

**Fig 2.**
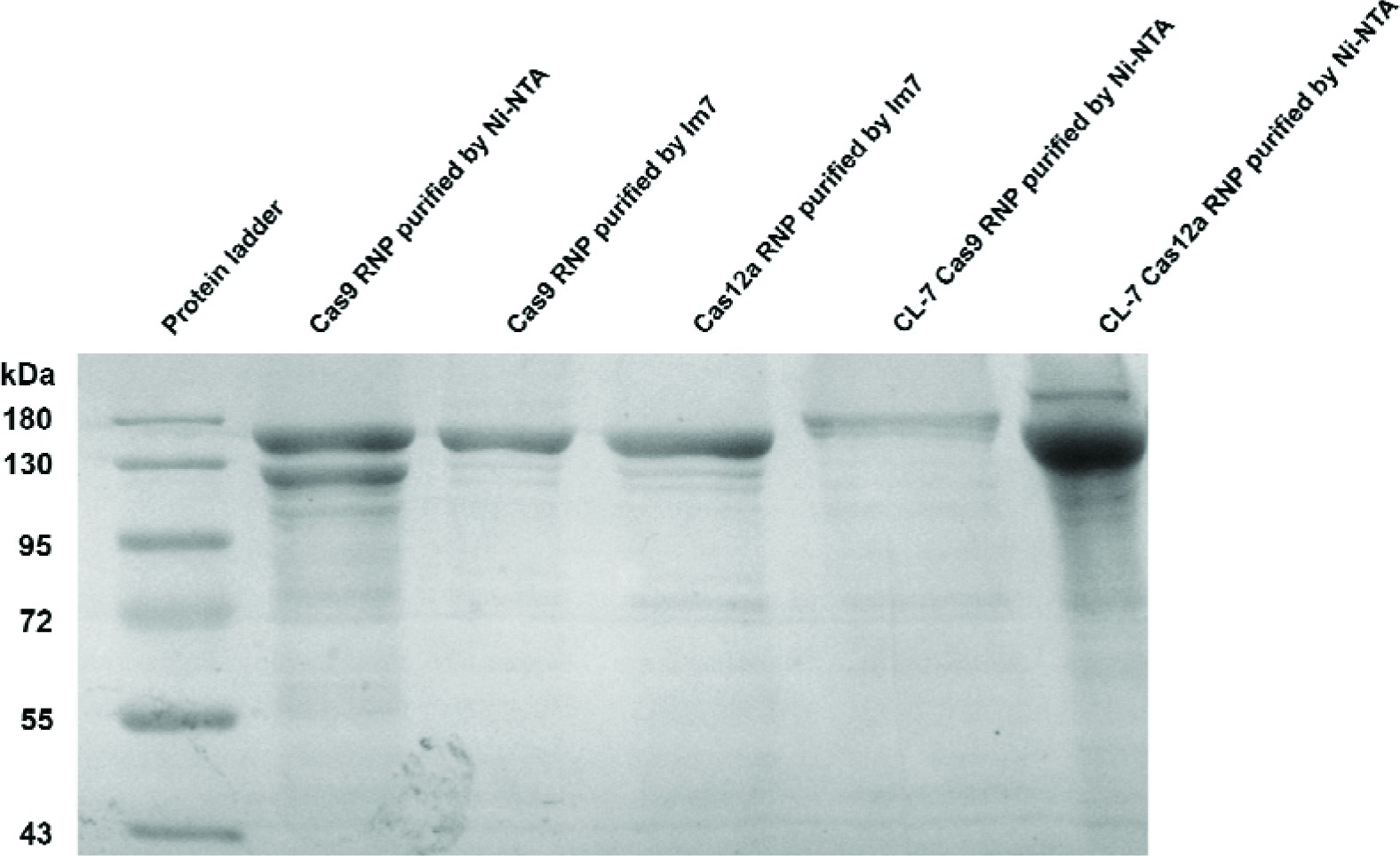
12% SDS-PAGE of purified Cas RNP by Ni-NTA or Im7 affinity columns.

The RNP prepared by the in vitro method currently used (Cho et al. 2013) is very unstable so that a sufficient amount of RNase inhibitors are required to prevent degradation of gRNA. By our co-expression method (Li et al. 2018), there is no needs to add RNase inhibitors in the whole purification and storage procedures. Herein, we found that the Cas9 or Cas12a RNP made by our in vivo method is also pretty stable that can maintain full nuclease activity when storing at −20°C for up to half a year.

### Characterization of the purified Cas RNP

To determine their in vitro nuclease activity, we observed that 300 ng plasmids (**Fig. 3a**) were cleaved in less than 30 minutes at 37°C when 200 ng of RNPs were added (SI). Accordingly, Cas9, CL7-Cas9 and Cas12a RNP can fully cleave the target plasmids. The endonuclease activity of our purified RNP is comparable to that of commercial Cas enzyme, indicating that the method can be industrially applicable to produce CRISPR-associated RNP.

**Fig 3.**
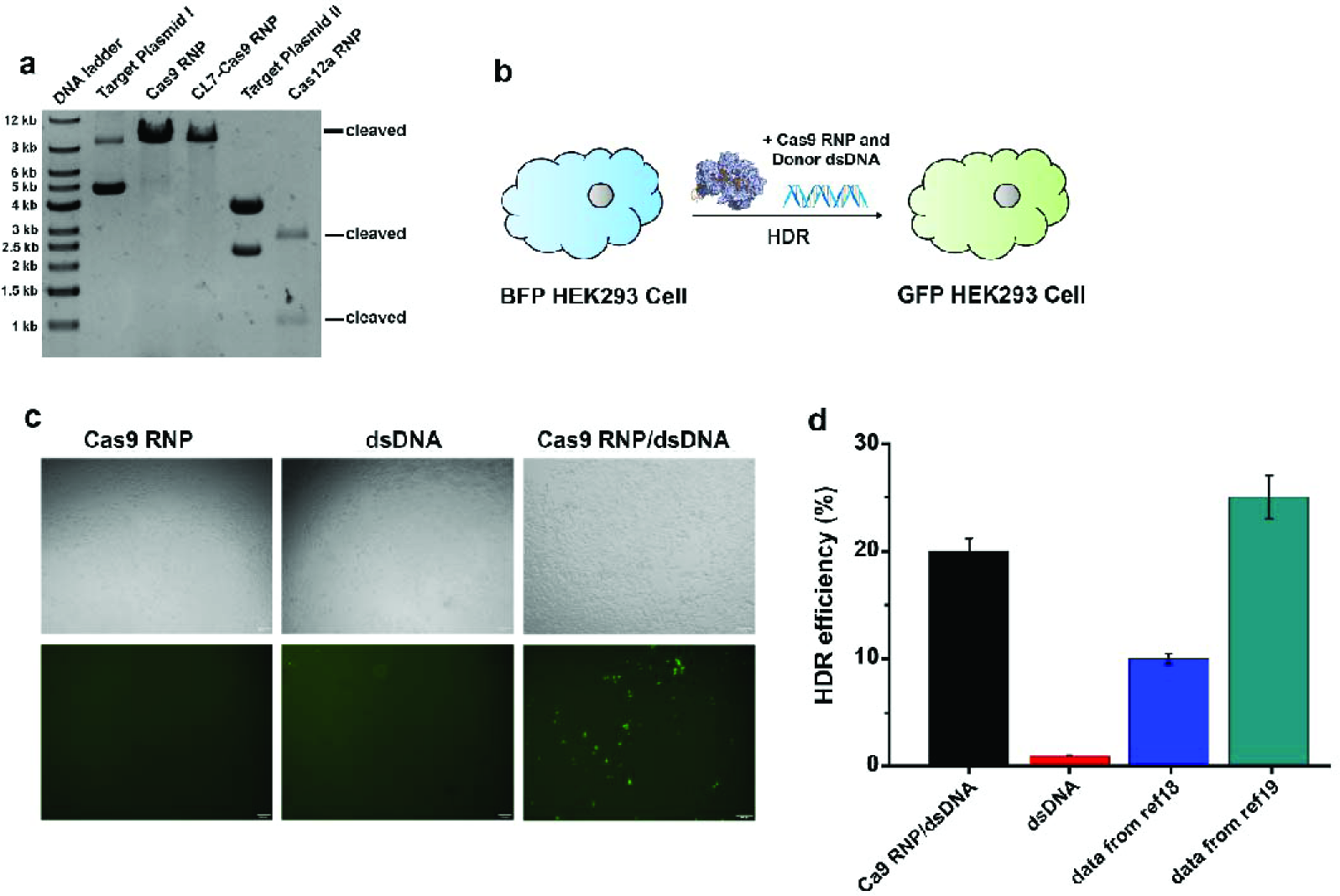
Endonuclease activity assays of purified Cas RNP. (a) In vitro cleavage of target plasmid I (a single cutting site) by Cas9 RNP or CL7-Cas9 RNP, as well as target plasmid II (two cutting sites) by Cas12a RNP. (b) Delivery of Cas9 RNP and donor dsDNA in BFP-HEK293 cells can induce HDR to transfer them into GFP-HEK 293 Cells. (c) The bright-filed and fluorescent images of BFP-HEK293 cells after delivering Cas9 RNP (left panel), donor dsDNA (middle panel) and Cas9 RNP together with donor dsDNA (right panel) by lipofectamine CRISPRMAX in 48 hours later. (d) HDR efficiency was determined by GFP expression due to BFP editing. In total, 20% efficiency was observed when using Cas9 RNP/dsDNA comparing to 1% efficiency with addition of dsDNA alone. The previous values in refence 18 and 19 were also present here, indicating that the Cas9 RNP prepared by our method has comparable HDR editing efficiency.

To illustrate the in vivo homology dependent repair (HDR) efficiency of purified Cas9 RNP, we constructed an engineered BFP-HEK293 cell line as the reporter system (DeWitt et al. 2017; Lee et al. 2017). In principle, the BFP-HEK293 cells would be converted to GFP-expressing cells after co-delivering Cas9 RNP and donor dsDNA by lipofectamine CRISPRMAX (**Fig. 3b**). The determined HDR efficiency of our purified Cas9 RNP is about 20%, which is comparable to the RNP made by in vitro as previously reported (**Fig. 3c**) (DeWitt et al. 2017; Lee et al. 2017).

## Discussion

Recently, direct delivery of pre-formed Cas9 RNP has been proven as a promising method for genome editing with great advantages (Cho et al. 2013; DeWitt et al. 2017; Staahl et al. 2017; Yu and Liang 2016). However, it is often time-consuming and expensive to prepare Cas9 RNP in sufficient amounts by the current method through roughly mixing Cas9 and gRNA in vitro. n conclusion, we utilize an easily removeable CL7 fusion tag to overexpress Cas9 or Cas12a RNP in vivo in E. coli, enabling significantly improved production of RNP. By harnessing the ultra-high-affinity CL7/Im7 purification system, we achieved one-step purification of pure Cas RNP with profound nuclease activity in vitro and in vivo, indicating that the method has great potentials to prepare CRISPR-associated nucleases at large-scale, low cost and short time.

## Acknowledgments

We thank T.A. Liu for kindly gifting the genes involving astaxanthin biosynthetic pathway. This research was supported by Key Technical Innovation Foundation of Hubei Province (NO.2017ACA174 to L.X. Ma) and National Foundation of Hubei Province (NO. 201700963 to Y. Liu).

